# A NYN domain protein directly interacts with DCP1 and is required for phyllotactic pattern in Arabidopsis

**DOI:** 10.1101/2021.10.01.462782

**Authors:** Marlene Schiaffini, Clara Chicois, Aude Pouclet, Tiphaine Chartier, Elodie Ubrig, Anthony Gobert, Hélène Zuber, Jérôme Mutterer, Johana Chicher, Lauriane Kuhn, Philippe Hammann, Dominique Gagliardi, Damien Garcia

## Abstract

In eukaryotes, general mRNA decay requires the decapping complex. The activity of this complex depends on its catalytic subunit, DCP2 and its interaction with decapping enhancers, including its main partner DCP1. Here, we report that in *Arabidopsis*, DCP1 also interacts with a NYN domain endoribonuclease, hence named DCP1-ASSOCIATED NYN ENDORIBONUCLEASE 1 (DNE1). Interestingly, we find DNE1 predominantly associated with DCP1 but not with DCP2 and reciprocally, suggesting the existence of two distinct protein complexes. We also show that the catalytic residues of DNE1 are required to repress the expression of mRNAs *in planta* upon transient expression. The overexpression of DNE1 in transgenic lines leads to growth defects and transcriptomic changes related to the one observed upon inactivation of the decapping complex. Finally, the combination of *dne1* and *dcp2* mutations, revealed a functional redundancy between DNE1 and DCP2 in controlling phyllotactic pattern formation in *Arabidopsis*. Our work identifies DNE1, a hitherto unknown DCP1 protein partner highly conserved in the plant kingdom and identifies its importance for developmental robustness.

**One-sentence summary:** DNE1, a NYN domain protein interacts with the decapping activator DCP1 and, together with DCP2, specify phyllotactic patterns in *Arabidopsis*.

## INTRODUCTION

Messenger RNA (mRNA) decay is crucial for the regulation of gene expression and is required for stress response and developmental transitions. Most mRNAs are degraded by the 5’-3’ pathway which requires decapping of mRNA (Sorenson et al., 2018; Tuck et al., 2020). mRNA decapping is a highly conserved mechanism in eukaryotes and consists in the hydrolysis of the 5’ m7G cap structure of the mRNA. Decapping produces an unprotected 5’ phosphate extremity, rapidly attacked by the cytosolic 5’-3’ exoribonuclease, XRN1 in mammals and XRN4 in plants. The catalytic subunit of the decapping complex is the Nudix hydrolase DCP2, which requires interactions with enhancers of decapping to switch from its inactive to its active form (Chang et al., 2014; Wurm and Sprangers, 2019). These activators notably include DCP1, LSM14A, EDC4, PAT1, the LSm1-7 complex and the DDX6 RNA helicases in metazoans (She et al., 2008; Arribas-Layton et al., 2013). mRNA decapping represents a limiting step of many cellular RNA decay pathways. Decapping notably operates after mRNA deadenylation (Couttet et al., 1997), downstream of the action of microRNAs (Nishihara et al., 2013), on mRNAs containing premature termination codons, through the action of Nonsense Mediated Decay (Lejeune et al., 2003), as well as on mRNAs with specific stem-loop structures in their UTRs in Staufen Mediated Decay (Kim and Maquat, 2019).

In plants, decapping has been mainly studied in *Arabidopsis thaliana*, whose genome encodes orthologs of DCP2, DCP1, LSM14A called DECAPPING 5 (DCP5), three DDX6-like RNA helicases called RH6, RH8 and RH12 as well as an ortholog of EDC4 called VARICOSE (VCS) and its presumed inactive homolog VARICOSE RELATED (VCR) (Deyholos et al., 2003; Xu et al., 2006; Xu and Chua, 2009; Chantarachot et al., 2020). Null mutants of *DCP1, DCP2, DCP5* and *VCS* show defects in the formation of the vasculature of embryonic leaves and are seedling lethal (Deyholos et al., 2003; Xu et al., 2006). A recent transcriptome-wide decay rate analysis in *Arabidopsis* revealed the major role of VCS in mRNA decay, with VCS involved in the decay of 68% of mRNAs (Sorenson et al., 2018). Although the importance of components of the decapping machinery was clearly established for its role in plant growth, biotic and abiotic stress response, only very limited information about the corresponding protein complexes was gathered to date (Xu and Chua, 2012; Soma et al., 2017; Yu et al., 2019; Kawa et al., 2020). The only data were obtained early on by *in vitro* tests and/or transient expression strategies, but in contrast to other organisms, was not yet approached by unbiased strategies designed to study protein complexes *in vivo*. Fundamental differences in the composition of this complex in the plant kingdom, including the existence of plant specific partners could thus not be assessed.

In this work, we used an unbiased approach coupling immunoprecipitations (IPs) and mass spectrometry to define the interactome of the decapping activator DCP1 and the decapping enzyme DCP2. In addition to characterize the *Arabidopsis* decapping complex, our work identifies a NYN domain endoribonuclease as a direct protein partner of the decapping activator DCP1, therefore named DCP1-ASSOCIATED NYN ENDORIBONUCLEASE 1 (DNE1). The closest homologue of DNE1 in mammals is Meiosis regulator and mRNA stability factor 1 (MARF1), a NYN domain endoribonuclease necessary for meiosis progression and retrotransposon surveillance in oocytes (Su et al., 2012a; Su et al., 2012b; Nishimura et al., 2018). MARF1 is associated to the core decapping complex formed by DCP1, DCP2 and EDC4 (Bloch et al., 2014). In contrast, we show here data suggesting the preferential association of DNE1 with DCP1 over DCP2. A phylogenetic analysis indicates that DNE1 is a highly conserved NYN domain protein in streptophytes and is likely an active enzyme as its catalytic residues are required to repress mRNAs upon transient expression. Furthermore, using a transcriptomic approach, we identified that overexpression of DNE1 leads to transcriptomic changes showing similarities with the ones observed upon inactivation of decapping. Finally, we identified that phyllotactic defects appear in *dne1 dcp2* double mutants demonstrating the redundant functions of DNE1 and DCP2 in developmental robustness.

## RESULTS

### Identification of proteins associated with the decapping complex in *Arabidopsis*

To determine the composition of the decapping complex in *Arabidopsis*, we used an unbiased proteomic approach. This approach is based on IP experiments coupled to mass spectrometry analysis using transgenic lines expressing either a functional yellow fluorescent protein (YFP) fusion of DCP1 expressed under its endogenous promoter or a functional version of DCP2 expressed under the 35S CaMV promoter (Goeres et al., 2007). Both IPs were efficient and provided access to both DCP1 and DCP2 enriched extracts (Fig. 1A, Supplementary Table S1). These extracts were then analyzed by liquid chromatography coupled to tandem mass spectrometry (nanoLC-MS/MS). Wild-type non-transgenic plants were used as controls for the enrichment analysis. This strategy allowed the identification of 15 proteins significantly enriched with DCP1 (Fig. 1B, Supplementary Table S2) and 8 proteins significantly enriched with DCP2 (Fig. 1C, Supplementary Table S3). The most enriched protein found with both DCP1 and DCP2 is VCS, which validates our approach. Of note, we also observed a strong enrichment of VCR, previously proposed not to be involved in decapping (Xu et al., 2006). Specifically enriched with DCP1, we found the DDX6-like RNA helicases RH6, RH8 and RH12, known to contact the decapping complex in mammals (Ayache et al., 2015) and that we recently found to co-purify with DCP5 and with the key player of NMD UPF1 (Chicois et al., 2018). An RNA helicase of the DDX3 family, RH52, in addition to 9 other proteins of unknown functions in RNA degradation was also specifically enriched. Among these factors, our attention was particularly drawn by a putative endonuclease (AT2G15560), enriched in DCP1, but not present in DCP2 IPs. We knew from our previous work on UPF1 interactome, that this protein also co-purified with both UPF1 and DCP5, and colocalized with UPF1 in cytoplasmic foci (Chicois et al., 2018). Because this predicted endonuclease was repeatedly found associated with RNA decay factors, we sought to further characterize its interactome, its influence on the transcriptome and its function in *Arabidopsis*. Based on the results presented hereafter, this protein was named DCP1-ASSOCIATED NYN ENDORIBONUCLEASE 1 (DNE1).

**Figure 1.**
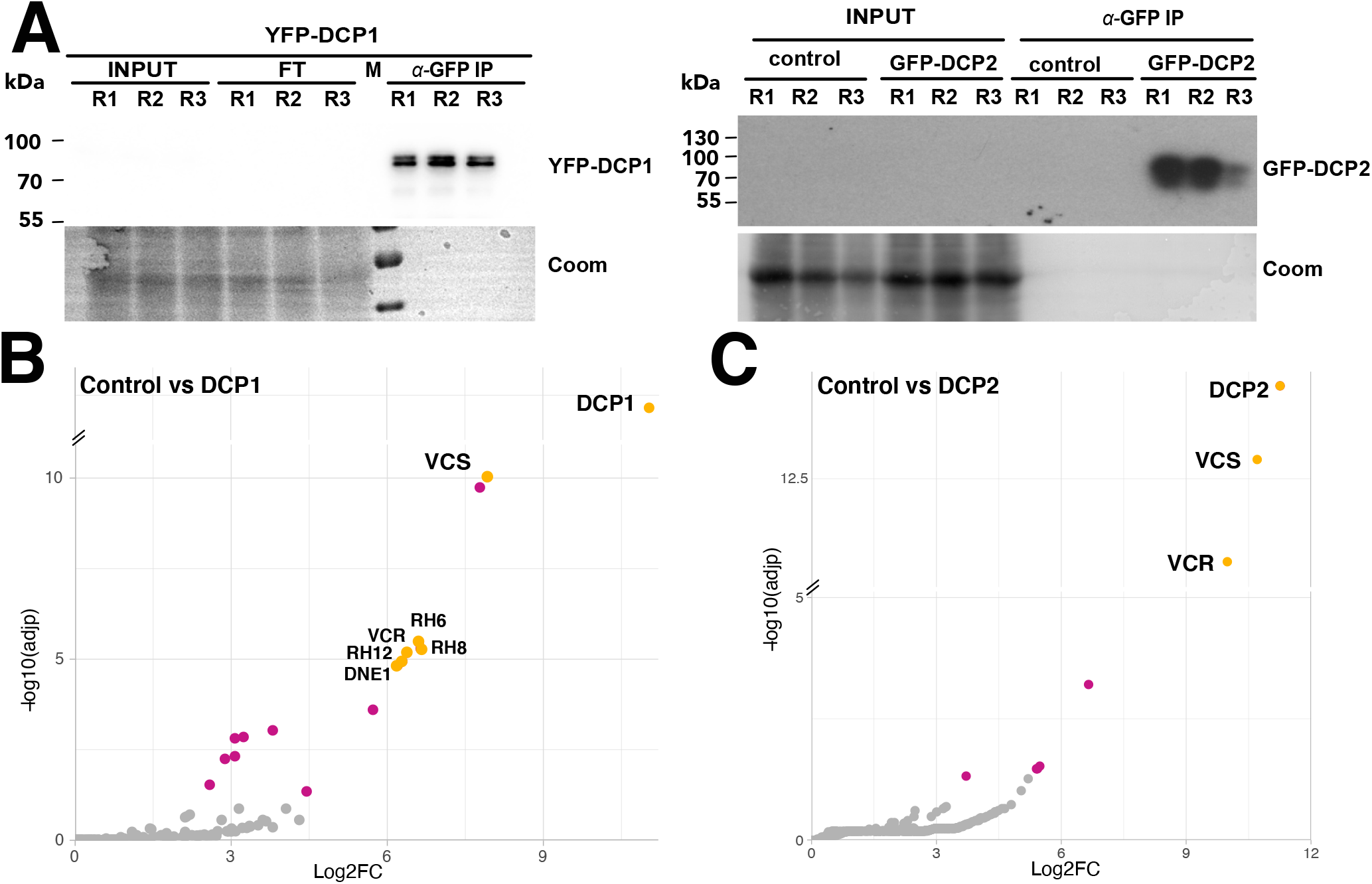
Identification of proteins associated with the decapping complex components DCP1 and DCP2. **(A)** Western blot analysis of GFP immunoprecipitates (IPs) performed in triplicate on extracts from YFP-DCP1 *dcp1-3* and from GFP-DCP2 *tdt-1* complemented lines. Wild-type plants used as negative controls are shown in the right panel (control). **(B)** Semi-volcano plot of proteins enriched in YFP-DCP1 IPs (n=6), results provided in Supplementary Table S2. **(C)** Semi-volcano plot of proteins enriched in GFP-DCP2 IPs (n=3), results provided in Supplementary Table S3. Control IPs (n=6) for results presented in B and C. Colored points (yellow and magenta) indicate proteins significantly enriched with Log FoldChange (Log2FC) > 1 and adjusted p-value (adjp) < 0.05. Yellow points highlight expected partners of the decapping complex and DNE1. Coomassie staining (Coom), protein ladder (M), flow-through (FT) and immunoprecipitated fractions (α-GFP IP).

### DNE1 directly interacts with DCP1

Transcriptomic data indicated that DNE1 is broadly expressed during plant development together with DCP1 and DCP2, albeit often at lower levels (Supplementary Fig. S1). As a first step in DNE1 characterization, we determined its interactome by IPs coupled to LC-MS/MS analysis. Previously designed transgenic lines expressing the endonuclease fused to GFP (Chicois et al., 2018) were used to perform DNE1 IPs (Fig. 2A). Four factors known to be associated with the decapping complex were among the proteins significantly enriched in DNE1 IPs: DCP1, VCS, VCR and UPF1 (Fig. 2B, Supplementary Table S4). Interestingly, DCP1 was the most enriched protein in DNE1 IPs. Considering that DNE1 was also among top partners in DCP1 IPs, we tested for a possible direct interaction between DCP1 and DNE1 by yeast two hybrid (Y2H) assays. Indeed, yeast expressing both DCP1 and DNE1 showed fully restored growth on selective yeast media (Fig. 2C). Growth of the Y2H strain was also restored by co-expressing DCP1 and a version of DNE1 mutated in a predicted catalytic residue (D153N), suggesting that this interaction is independent of its catalytic activity.

**Figure 2.**
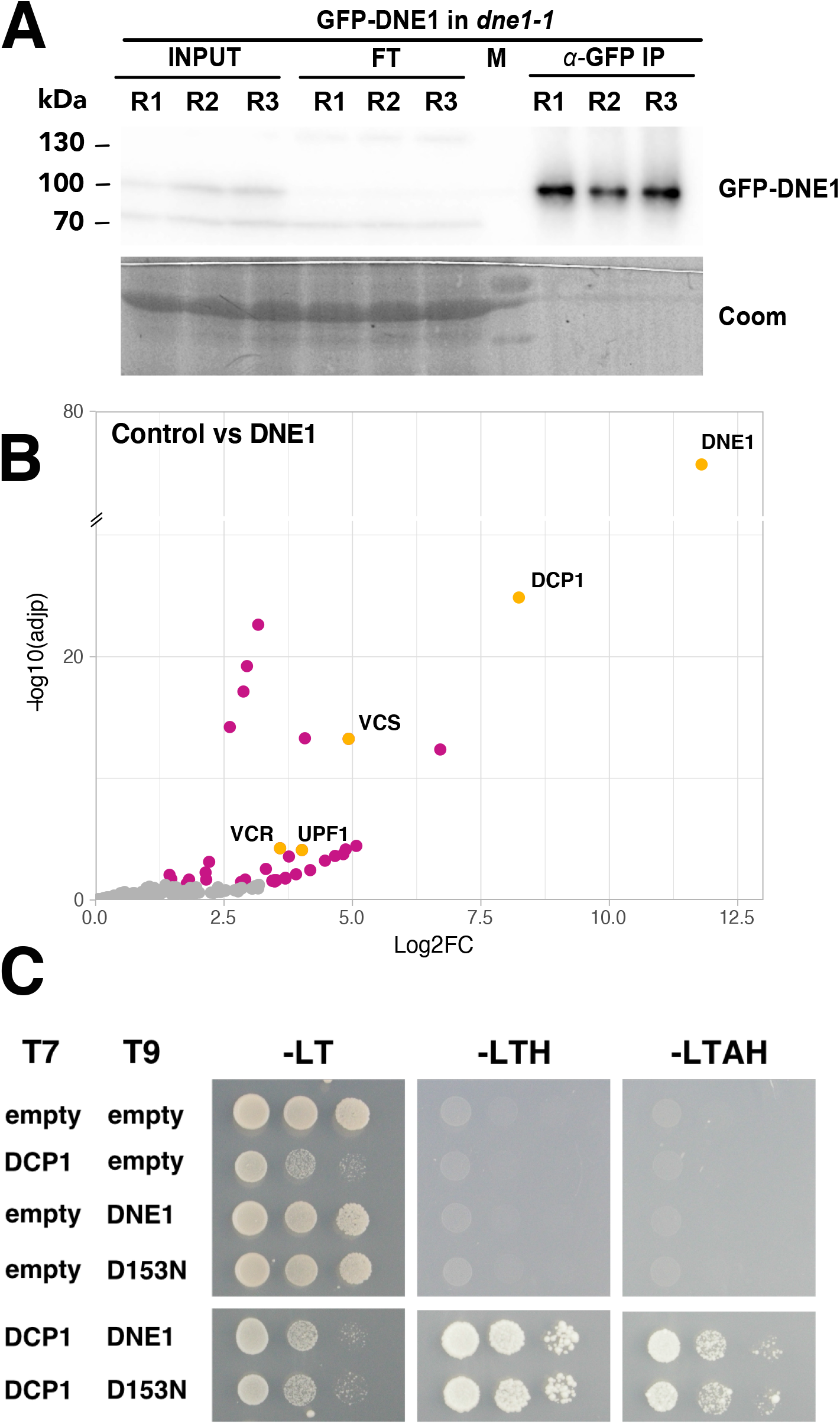
Identification of proteins associated with the DCP1-associated endonuclease DNE1. **(A)** Western blot analysis of GFP IPs performed in triplicate on extracts from GFP-DNE1 *dne1-1* lines. **(B)** Semi-volcano plot of proteins enriched in GFP-DNE1 IPs (n=8), control IPs (n=9), results provided in Supplementary Table S4. The volcano plot is represented as in Figure 1. **(C)** Specific growth on selective media for the DCP1-DNE1 and DCP1-D153N combinations highlights the direct interaction between DCP1 and DNE1. Minimal SD medium –LT, -LTH and –LTAH were used, in which Adenine (A); Histidine (H); Leucin (L); Tryptophan (T). 5mM 3-AT was used to avoid autoactivation. T7: pGADT7 AD (LEU2); T9: pGBT9 BD (TRP1).

To our surprise, DCP2 was not significantly enriched in neither DNE1 (Fig. 2B) nor DCP1 IPs (Fig. 1B), and only a few mRNA decapping proteins were present in the DCP2 IPs (Fig. 1C). To test if the non-detection of DCP2 with DCP1 and the low efficiency of the DCP2 IPs can be solved by stabilizing transient or weak interactions, DCP1, DCP2 and DNE1 IPs were repeated with formaldehyde crosslinked protein extracts. In these conditions, we observed the association between DCP1 and DCP2 in both DCP1 and DCP2 IPs (Fig. 3A, 3B; Supplementary Tables S5, S6). In addition, we also found additional co-purifying partners linked to decapping in DCP2 IPs (including DCP1, RH6, RH8, RH12, UPF1 and PAT1, Fig. 3B; Supplementary Tables S6). Interestingly, DNE1 was only detected together with DCP1 but was again absent from DCP2 purifications (Fig. 3). In contrast, DCP1 was found together with both DCP2 and DNE1 (Fig. 3B, 3C; Supplementary Tables S6, S7). These data obtained upon crosslinking solve the initial low efficiency of DCP2 IPs. Although no definitive conclusions can be drawn from negative results in IP experiments, these data taken together are coherent with the existence of two different complexes, one comprising DCP1-DCP2 and one containing DCP1-DNE1.

**Figure 3.**
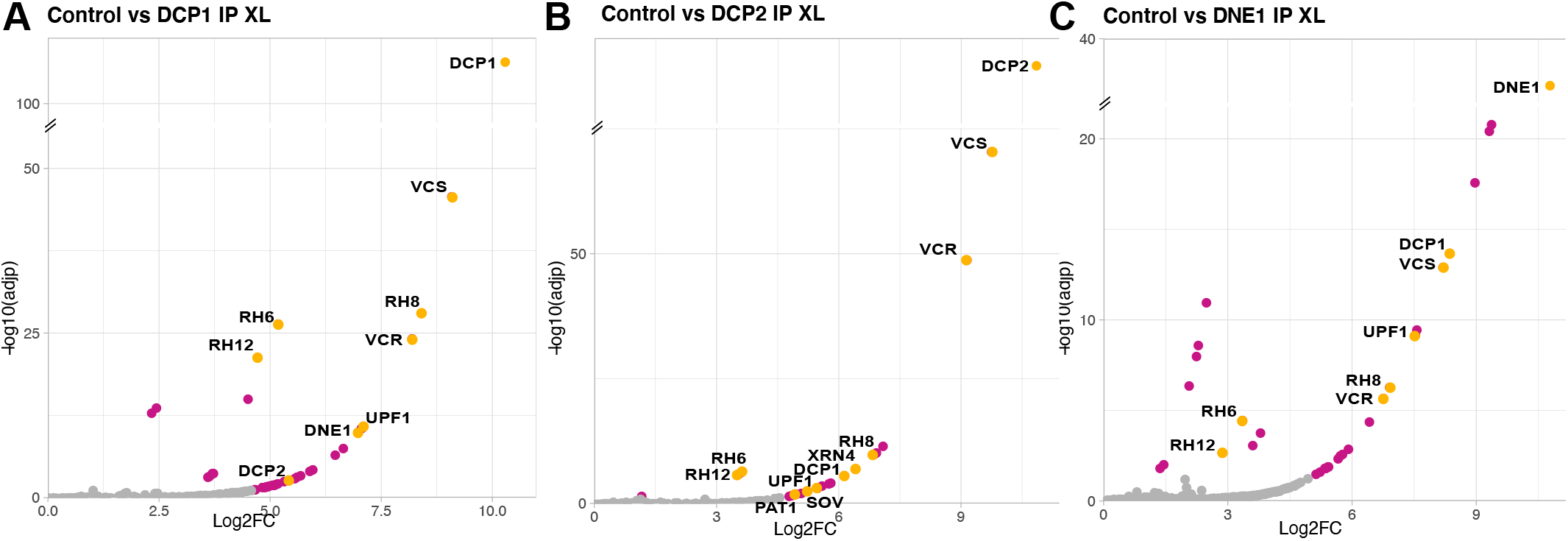
Crosslinked immuno-precipitations improve the sensitivity for the identification of proteins associated with DCP1, DCP2 and DNE1. **(A)** Semi-volcano plot of proteins enriched in YFP-DCP1 crosslinked IPs (n=4), results provided in Supplementary Table S5. **(B)** Semi-volcano plot of proteins enriched in GFP-DCP2 crosslinked IPs (n=4), results provided in Supplementary Table S6. **(C)** Semi-volcano plot of proteins enriched in GFP-DNE1 crosslinked IPs (n=4), results provided in Supplementary Table S7. Control IPs (n=4) for results presented in A, B and C. Colored points (yellow and magenta) indicate proteins significantly enriched with Log FoldChange (Log2FC) > 1 and adjusted p-value (adjp) < 0.05. Yellow points highlight expected partners of the decapping complex and DNE1, cytosolic exoribonucleases XRN4 and SOV and the NMD protein UPF1.

### DNE1 is a component of P-bodies

Our previous work localized DNE1 in cytoplasmic foci with UPF1 upon transient expression, identifying DNE1 as a putative novel component of P-bodies (Chicois et al., 2018). To further validate this localization of DNE1 in P-bodies, we produced transgenic *Arabidopsis* lines expressing DNE1 (fused either to GFP or RFP) together with archetypal markers of either stress granules (PAB2-RFP) or P-bodies (YFP-DCP1, UPF1-RFP). In these lines, DNE1 did not colocalize with PAB2, both in the absence of stress, when PAB2 exhibits a diffuse localization (Pearson correlation coefficient 0.18, Fig. 4) and under heat stress, when PAB2 is relocated in stress granules (Pearson correlation coefficient 0.35; Fig. 4). By contrast, DNE1 perfectly colocalized with its protein partner DCP1 and with UPF1 in P-bodies (Pearson correlation coefficient 0.9, Fig. 4). We therefore concluded that DNE1 is a *bona fide* component of P-bodies in *Arabidopsis*.

**Figure 4.**
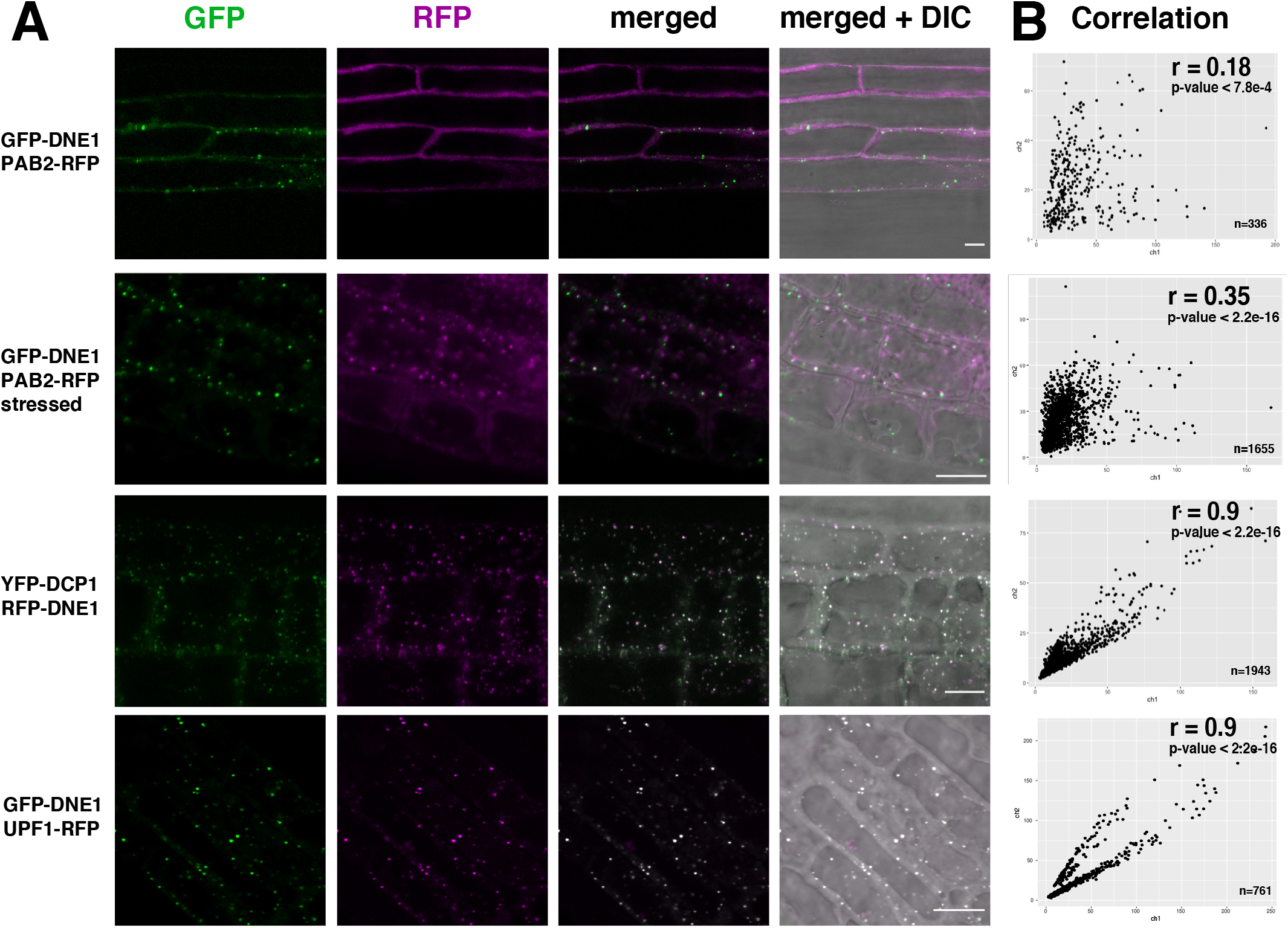
DNE1 co-localizes with DCP1 and UPF1 in p-bodies. **(A)** Confocal microscopy co-localization study of DNE1 with the stress granule marker PAB2-RFP, the P-body markers YFP-DCP1and UPF1-RFP in stable *Arabidopsis* transformants. A 30 min heat stress at 37°C was applied to GFP-DNE1 PAB2-RFP to induce stress granule formation. Scale bar: 10µm. **(B)** Dot plot showing the quantification of foci co-localization in the green (ch1, y-axis) and red (ch2, x-axis) channels; The number of foci analyzed (n) is indicated on the plot. The calculated Pearson’s correlation coefficient (r) and p-values are indicated.

### DNE1 is a well conserved NYN domain protein in streptophytes and composed of three domains

To determine the evolutionary conservation of DNE1, we looked for orthologues of the *Arabidopsis* DNE1 protein in major plant lineages (rhodophytes, chlorophytes and streptophytes). We refined the results based on the presence of the structural domains found in *Arabidopsis* DNE1 that are one NYN domain and two consecutive OST-HTH domains. We found clear orthologues of DNE1 in basal streptophytes algae, mosses, ferns, gymnosperms and all flowering plants (Fig. 5A). We could not find NYN domains associated with predicted OST-HTH domains in the rhodophytes and chlorophytes phyla. However, we do not exclude the hypothesis that OST-HTH sequences could be more degenerated from the consensus in these lineages and therefore not recognized by the prediction software. Indeed, the sequence of NYN domain is more conserved (identity and similarity) than the OST-HTH domains in the streptophytes, especially the second OST-HTH (Supplementary Table S8). Of note, the second OST-HTH domain was not found in *Klebsormidium* a basal streptophytes alga but KfDNE1 was the most related enzyme to AtDNE1 in this specie. Overall, DNE1 is conserved in most if not all species of the streptophyte lineage indicating that its main function in plants is likely under selection pressure.

**Figure 5.**
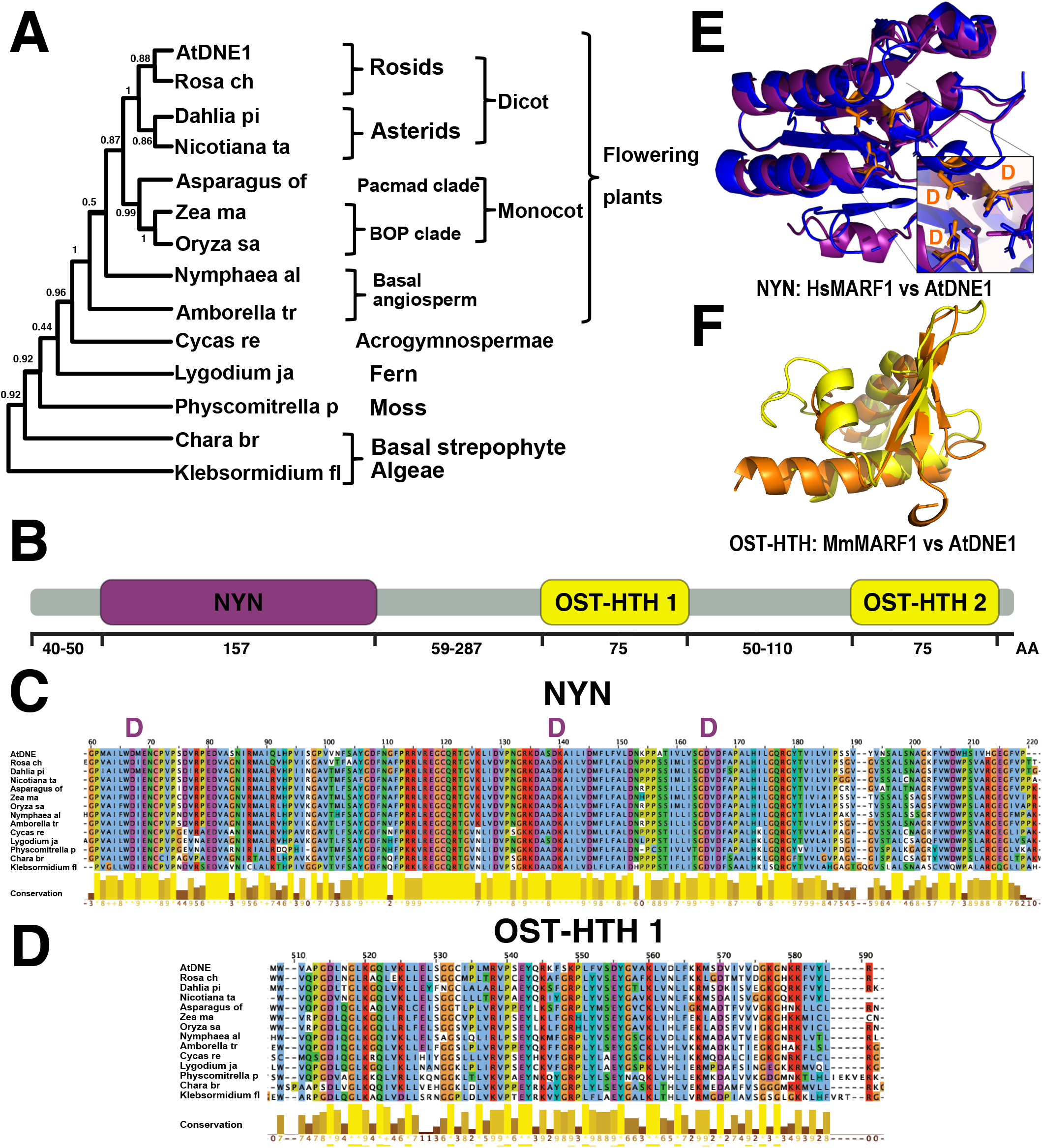
DNE1 is an evolutionary conserved NYN domain protein harboring two OST-HTH modules. **(A)** DNE1 phylogenetic tree obtained with the maximum likelihood method. Bootstrap values are indicated for each node. **(B)** Schematic domain structure of DNE1. In purple the catalytic NYN domain, in orange the OST-HTH predicted RNA binding domains. **(C)** Multiple alignment of amino-acid sequences of NYN domains from DNE1 plant orthologs as in (A). In purple the conserved aspartic acid residues important for catalysis. **(D)** Multiple alignments of amino-acid sequences of OST-HTH 1 domains from DNE1 plant orthologs as in (A). **(E)** Structural alignment of the predicted tridimensional structure of AtDNE1 NYN domain sequence (in purple) with the tridimensional crystal structure of *Hs*MARF1 NYN domain [6fdl, (Nishimura et al. 2018), in blue]. Conserved D residues are shown in orange. **(F)** Structural alignment of the predicted tridimensional structure of AtDNE1 OST-HTH1 domain sequence (in yellow) with the tridimensional crystal structure of *Mm*MARF1 OST-HTH1 domain [5yad, (Yao et al., 2018), in orange].

The typical structure of DNE1 is characterized by the presence of a NYN domain in the N-terminal part of the protein and of two OST-HTH domains in the C-terminal part (Fig. 5B). The length between the NYN domain and the two OST-HTH domains is variable with a larger region in early divergent species compared to more recent species (Fig. 5B). The most compact enzymes are found in the Brassicaceae (*Arabidopsis* family) with proteins whose sizes are under 500 amino-acids compared to 600 aa in average for conifers and more than 700 aa in *Klebsormidium*. The alignment of the NYN and the OST-HTH1 domains shows that the catalytic domain is extremely conserved at the amino acid level, feature supported by the identity and similarity chart (Supplementary Table S8) while the RNA binding domains are slightly less conserved (Fig. 5C, 5D, Supplementary Table S8). Importantly, we identified the three aspartic acid residues known to be necessary for the enzymatic activity of NYN domains (Fig. 5C, 5E). These residues are strictly conserved in all plant species assessed (Fig. 5C). The closest homologue of DNE1 in mammals is the endoribonuclease MARF1. In mammals, MARF1 functions in female germlines where it is crucial for meiosis and defenses against damages caused by retrotransposons to the oocyte’s genome (Su et al., 2012a; Su et al., 2012b). The tridimensional structure predictions of DNE1 protein domains fit well into the known tridimensional crystal structures of MARF1 domains, both for the NYN (Fig. 5E) and for the OST-HTH (Fig. 5F) domains. In addition, the conserved Asp catalytic residues (in orange Fig. 5E) share a similar location in the structural alignment of MARF1 and DNE1 NYN domains that strongly suggests that DNE1 is also an active enzyme. This possibility was supported by *in vivo* transient expression experiments of WT or a catalytic mutant of DNE1 (D153N) in *N. benthamiana*. In this experiment either RFP-DNE1 or RFP-DNE1^D153N^ catalytic mutant was co-expressed with a reporter *GFP* mRNA in *N. benthamiana* leaves. A dramatic drop in the accumulation of the co-expressed *GFP* mRNA and three endogenous mRNAs was observed for three independent replicates 2.5 days after leaf infiltration of only RFP-DNE1 but not RFP-DNE1^D153N^ (Fig. 6). Of note, this difference is not due to a lower expression of the DNE1-D153N mutant as RFP-DNE1-D153N accumulates to higher levels that RFP-DNE1 in this experimental setup (Supplementary Fig. S2). This result indicates that the transient expression of DNE1 represses the accumulation of mRNAs and that this repression requires a wild-type NYN domain, supporting that DNE1 is an active NYN domain enzyme.

**Figure 6.**
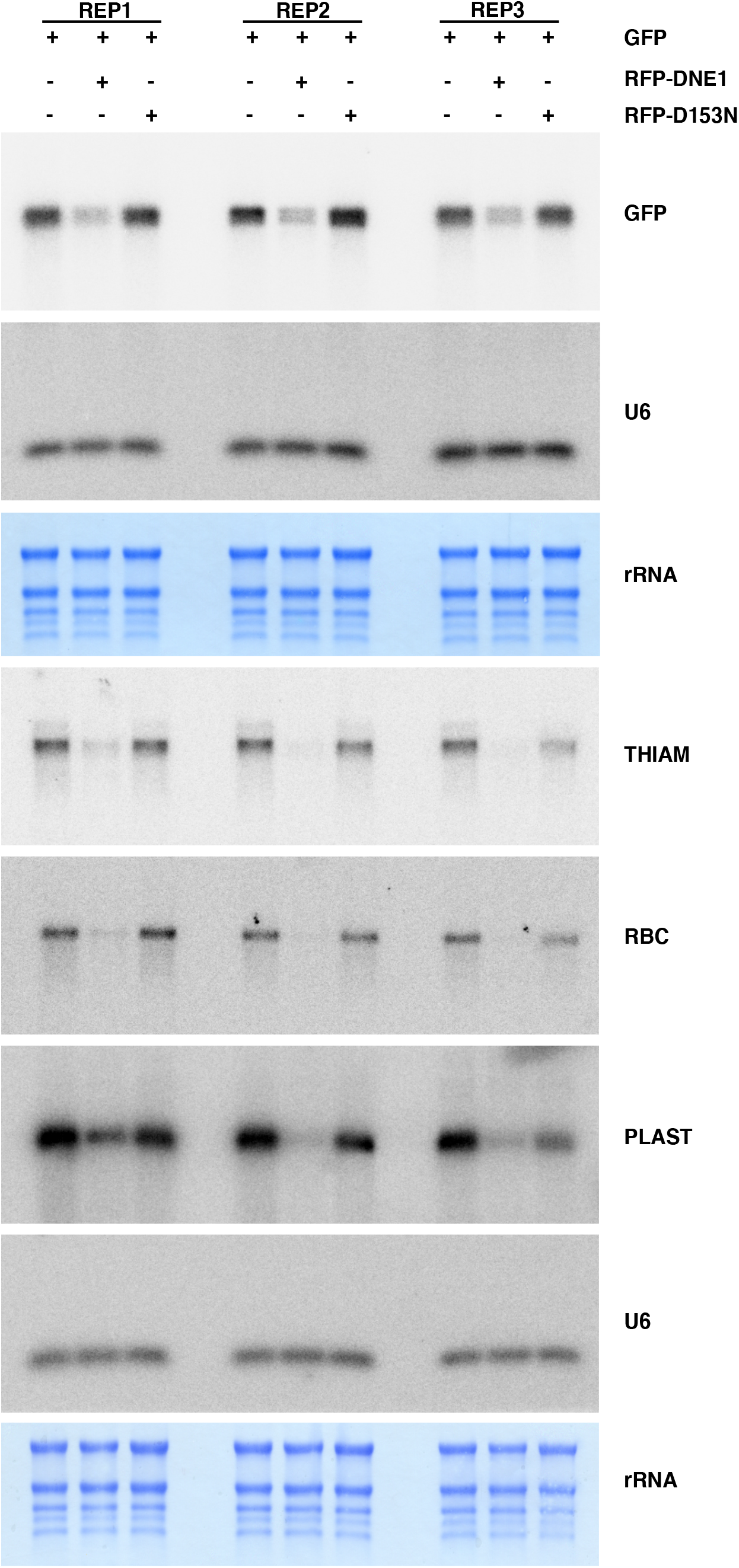
Transient expression of DNE1 impairs the expression of a co-expressed and endogenous mRNAs. Northern blot analysis showing the impact of the co-expression of DNE1 WT (WT) or DNE1 D153N (D153N) on mRNA accumulation in *N. benthamiana*, empty plasmid (-) is used as a control. Methylene blue staining showing ribosomal RNAs (rRNA) and a U6 probe are used as loading controls.

### DNE1 overexpression perturbs plant growth and show similar changes in gene expression than decapping mutants

To analyze the influence of DNE1 on the transcriptome, we tested the impact of the deregulation of DNE1 on *Arabidopsis* transcriptome using RNA-seq. For this purpose, we used three different genetic backgrounds: an insertion mutant (*dne1-1*), which harbors a T-DNA insertion within the *DNE1* coding sequence, and transgenic lines overexpressing protein fusions with GFP of either a WT (DNE1 OE) or a DNE1 point mutant (D153N OE) in *dne1-1* background. The point mutation is in a codon coding for a conserved aspartic acid residue and predicted to abolish the catalytic activity of DNE1. Visual inspections made during the growth of *dne1-1* mutant seedlings did not reveal any morphological defects. In contrast, the overexpressing lines of both DNE1 and D153N both showed growth retardation (Fig. 7A, Supplementary Fig. S3 and S4). We selected homozygous lines expressing DNE1 and D153N. Of note, we used a D153N OE line expressing lower protein levels than the wild-type DNE1 OE lines (Fig. 7B) as D153N lines with stronger protein expression could not be propagated. We analyzed the transcriptome of these lines and compared them with the transcriptome of wild-type plants and the null decapping mutant *vcs-6*. The global differences in gene expression were visualized on a histogram (Fig. 7C). Very little variation in gene expression was observed in *dne1-1*, with only 8 significantly deregulated genes. In contrast, DNE1 OE and D153N OE lines showed substantial transcriptome changes, while as expected massive deregulations occurred in *vcs-6*. Both DNE1 OE and D153N OE lines showed similar gene deregulation trend with more upregulated than downregulated transcripts. Indeed, we observed that 93% and 77% of the significantly deregulated transcripts were upregulated in WDNE1 OE and D153N OE, respectively. This result suggests that the overexpression of wild-type or mutant DNE1 might have similar effect and that this effect is likely not solely due to catalytic activity. Importantly, D153N expression affected more broadly the transcriptome than the expression of DNE1, with *ca* ten times more deregulated genes in D153N compared to DNE1. We found 64 commonly upregulated genes in DNE1 OE and D153N OE (Fig. 7D). The significance of the overlap between DNE1 OE and D153N OE deregulated genes was tested using a hypergeometric test. With P-values of 4.6 × 10^−42^ for upregulated genes and 4.1 × 10^−7^ for downregulated genes, this result indicates a similar signature of transcriptomic changes in DNE1 and D153N (Table S9).

**Figure 7.**
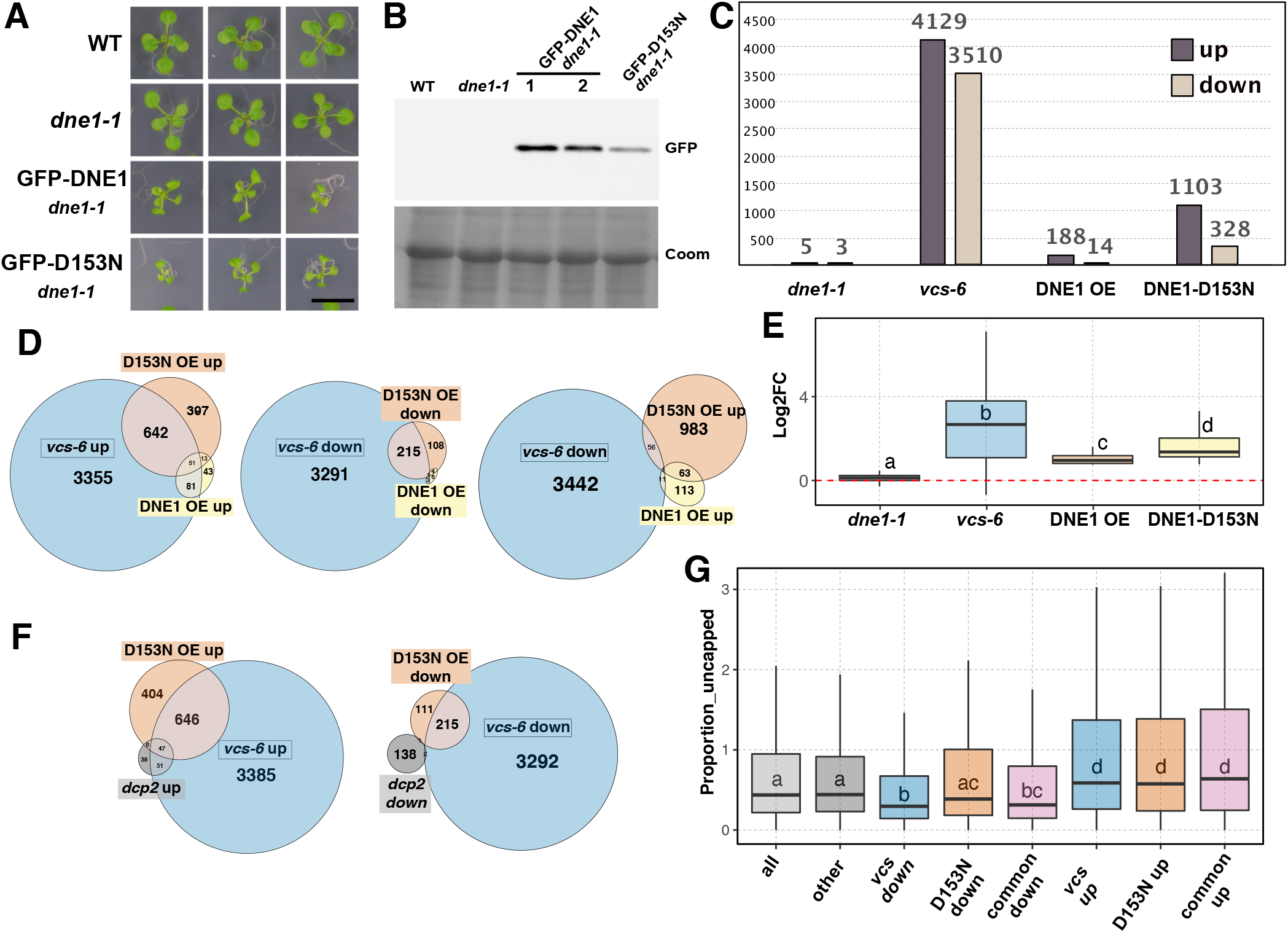
Altering DNE1 expression impairs plant growth and leads to the same gene deregulation signature than mutations in VCS and DCP2. **(A)** Pictures of WT, *dne1-1*, GFP-DNE1 and GFP-D153N overexpressors (OE) 14 days after planting. Scale bar: 1cm. **(B)** Analysis of GFP-DNE1 and GFP-D153N transgenic lines by Western blot using anti GFP antibody. Two independent plant lines 1 and 2 are analyzed for GFP-DNE1 and one for the GFP-D153N. Coomassie staining used as a loading control (Coom). **(C)** Histogram showing the global changes in gene expression based on RNAseq analyses in *dne1-1*, GFP-DNE1 OE, GFP-D153N OE and *vcs-6* compared to WT. Genes were considered as upregulated and downregulated when adjusted p-value<0.05 and Log2FC>0.75 or Log2FC<-0.75, respectively (n≥3). Y-axis represent the number of deregulated genes **(D)** Venn diagrams showing comparisons between significantly upregulated or downregulated genes in DNE1 OE or D153N OE lines and *vcs-6*. **(E)** Boxplot comparing the change levels (Log2FC) between the different genotypes of the 64 genes that are commonly upregulated in both DNE1 and D153N lines. **(F)** Venn diagrams showing comparisons between significantly upregulated or downregulated genes in D153N OE, *vcs-6* and the weak *dcp2* mutant *tdt-1*. **(G)** Box plot showing the proportion of uncapped transcripts as found by GMUCT in (Anderson et al., 2018) for all genes detected in RNAseq (all), genes not deregulated in any genotypes (other) and genes deregulated in *vcs-6* or D153N OE lines. Letters in (E) and (G) show statistically different groups based on a Wilcoxon rank-sum test.

We then tested whether gene deregulation in DNE1 and D153N OE lines was related to genes deregulated upon inactivation of decapping. We thus compared the list of genes deregulated in the two DNE1 OE lines with those deregulated in the null decapping mutant *vcs-6*. Strikingly we observed a significant overlap of genes upregulated in *vcs-6* and genes upregulated either in DNE1 OE or in D153N OE (P-values: DNE1 = 4.9 × 10^−66^, D153N = 2.8 × 10^−307^): 70% (132/188) and 63% (693/1103) of genes upregulated in DNE1 OE or in D153N OE respectively, are also up regulated in *vcs-6* (Fig. 7D left panel, Table S9). In contrast, and as a negative control, the comparison *vcs* downregulated versus DNE1 OE or D153N OE upregulated, gave no significant overlap with p-values of 1 (Fig. 7D right panel, Table S9). These results were validated by a Q-PCR experiment for a selection of 19 targets with different profiles (Supplementary Fig. S5, Supplementary Table S10). This assay validates our global analysis with a different experimental setup and strengthen the previous observations. Among the 13 genes validated as upregulated in DNE1 OE and/or in D153N OE tested, 11 loci were validated to be also upregulated in *vcs-6*, reinforcing the link between the overexpression of DNE1 and the effect of a mutation in VCS. In this analysis, 9 out of 13 upregulated genes were found more deregulated in D153N than DNE1, confirming the stronger effect of D153N expression compared to DNE1. Collectively those results indicate that overexpression of DNE1 altered gene expression in a manner reminiscent of *vcs* mutant plants, suggesting that DNE1, when overexpressed, alters the action of the decapping complex.

### Altering DNE1 expression or DCP2 action affects similar loci

As VCS is associated with DCP1, DCP2 and DNE1, and in order to test a possible functional link between the decapping enzyme DCP2 and DNE1, we then compared transcriptomic data obtained by microarray for *tdt-1*, a previously reported *dcp2* allele (Goeres et al., 2007), with our RNA-seq data of *vcs-6* and D153N OE. As expected, a significant overlap was found between the *vcs* and the *dcp2* upregulated genes (P-value 2.1 × 10^−37^) with 67.8% of genes upregulated in *dcp2* also upregulated in *vcs* (98/142, Fig. 7F). Interestingly, there was also a significant overlap of genes upregulated in *dcp2* and D153N (P-value of 3.1 × 10^−48^) with 37% of genes upregulated in *dcp2* also upregulated in D153N upregulated genes (53/142; Fig. 7F). Altogether, these analyses indicate a strong functional links between DNE1, VCS and DCP2. This comparison shows that the overexpression of D153N has perturbed the accumulation of a set of mRNAs also affected by mutation in *DCP2*. In support of this observation, we tested if mRNAs deregulated in D153N OE are enriched in targets of the decapping complex. We thus compared our RNA seq data for D153N OE and *vcs* with degradome sequencing data obtained by GMUCT (Anderson et al., 2018), a sequencing protocol which quantifies the accumulation of uncapped transcripts. Strikingly, in this comparison, only the upregulated genes in *vcs* and DNE1 D153N OE were enriched in loci accumulating uncapped transcripts (Fig. 7G). This results further strengthen our previous conclusion and validate that the deregulations observed in D153N OE seems to preferentially affect targets of the decapping complex.

### Mutations in DNE1 affect the robustness of phyllotaxis when DCP2 function is altered

To investigate the functional link between DNE1 and DCP2, we produced double mutants between *its1*, a weak allele of *dcp2* mutated in its catalytic domain (called *dcp2*^*its1*^ thereafter) and two *dne1* point mutants generated by CRISPR/Cas9 (*dne1-2* and *dne1-3*, Supplementary Table S11). The *dne1-3* mutant contains a 39 nt insertion leading a premature termination codon early in *DNE1* NYN domain, whereas *dne1-2* contains a 21nt in frame deletion at position 193, which predicts the production of a DNE1 protein with a 7 amino acids (aa) deletion in the catalytic domain. Neither the weak *dcp2*^*its*1^ mutant, nor the *dne1-2* and *dne1-3* showed any obvious morphological defects. In contrast, *dne1 dcp2*^*its*1^ double mutants showed defects in the maintenance of the phyllotactic pattern produced by the floral meristem (Fig. 8A). We quantified this defect by measuring divergent angles between successive siliques on the main stem (Fig. 8B). Overall, we made measurements for between 30 to 40 plants per genotype in three to four biological replicates, and measured divergent angles between 35 siliques per plant on the main stem. These measures clearly show an increased proportion of non-canonical divergent angles occurring specifically and reproducibly in *dne1 dcp2*^*its*1^ double mutants and not in *dcp2*^*its*1^ and *dne1* single mutants compared to WT. The defects observed in the double mutants were confirmed by the analysis of the divergent angles distribution using the Kolmogorov-Smirnov test, with p-values showing significant difference from the WT for *dcp2*^*its*1^ *dne1-3* (P-value 6.5 × 10^−8^) and *dcp2*^*its*1^ *dne1-2* (P-value 2.2 × 10^−16^, Fig. 8B, Table S12). These defects also appeared reproducibly on every biological replicate, when analyzed separately (Supplementary Fig. S6). This analysis demonstrates the synergistic effect of *dne1* and *dcp2*^*its*1^, revealing the redundant function of DCP2 and DNE1 in the establishment of phyllotaxis in the floral meristem. Interestingly, we could identify similar defects in *xrn4*, a mutant affected in the 5’-3’ mRNA degradation pathway (P-value 2.2 × 10^−16^, Fig. 8), suggesting that the observed phenotype results from an increased defect in mRNA degradation in the *dne1 dcp2*^*its*1^ double mutant.

**Figure 8.**
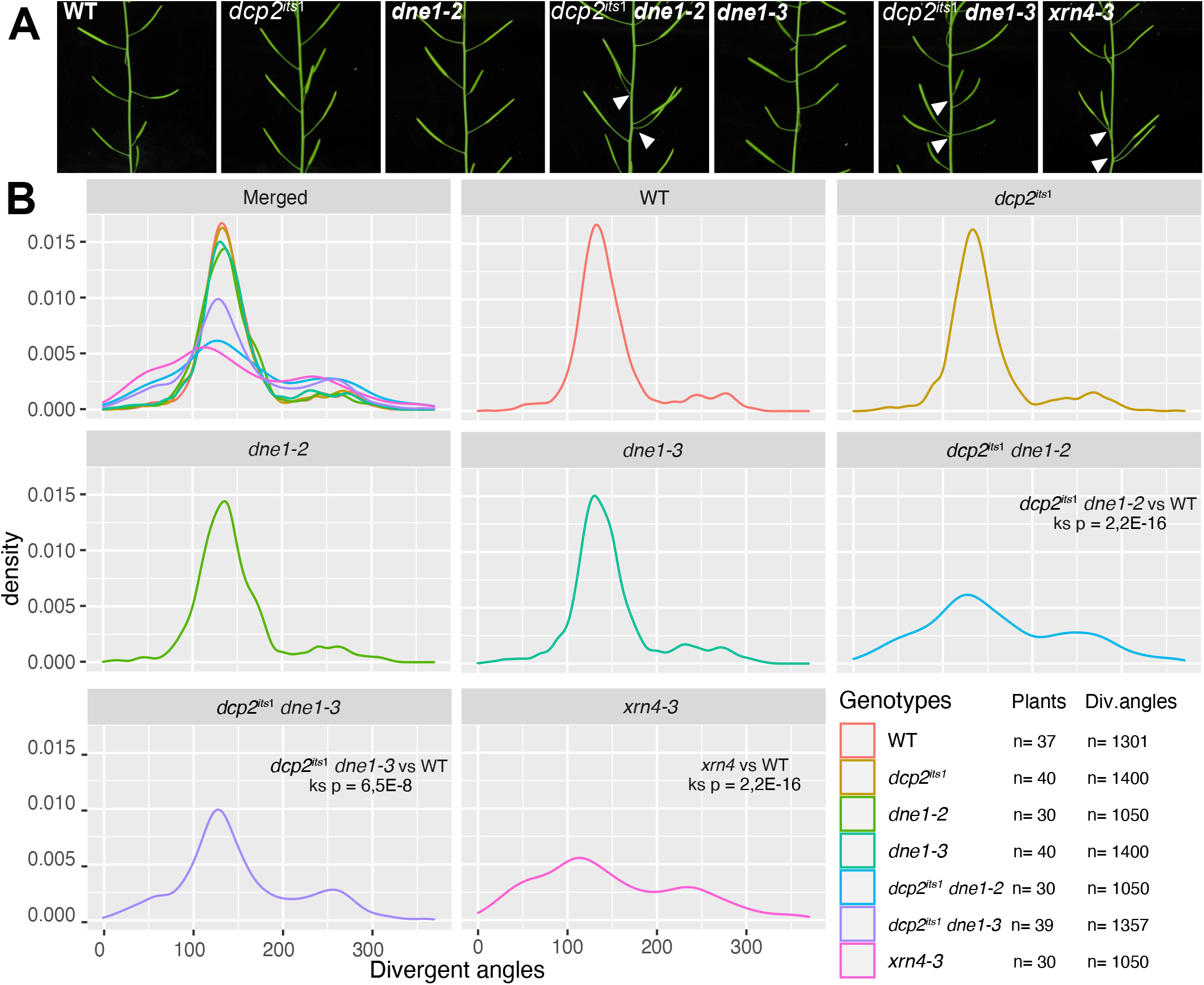
Synergistic effect of mutations in *dne1* and *dcp2* on phyllotactic pattern. **(A)** Pictures showing representative stems from the WT, *dcp2*^*its1*^, *dne1-2, dne1-2 dcp2*^*its1*^, *dne1-3, dne1-3 dcp2*^*its1*^ and *xrn4-3* plants. **(B)** Density plots showing the quantification of divergent angles from the genotypes shown in A. The analysis was performed on 3 to 4 biological replicates. Differences between divergent angles distribution was assessed using the Kolmogorov-Smirnov test, complete results are shown in Supplementary Table S12. The analysis is shown for each biological replicates separately in Supplementary Fig S6.

## DISCUSSION

In this study, we identified DNE1, a previously uncharacterized protein harboring an endoribonuclease domain of the NYN family and two OST-HTH domains, as a direct protein partner of DCP1. DNE1 accumulates together with DCP1 in P-bodies and is expressed throughout the plant lifecycle. We found that DNE1 is ubiquitous in streptophytes. This conservation for more than half a billion years suggests an important function. The perfect conservation in plants of aspartates residues known to be crucial for the catalytic activities of NYN-domain enzymes, strongly suggests that the activity of DNE1 is under a significant selection pressure (Matelska et al., 2017). The *in vivo* transient expression of wild-type DNE1 represses the expression of several mRNAs, whereas a catalytic mutant cannot, suggesting that DNE1 catalytic activity targets mRNAs. The constitutive overexpression of DNE1 in stable lines leads to growth defects and a similar gene deregulation signature than plants in which decapping is compromised. Finally, we present genetic evidences of a functional redundancy between DNE1 and DCP2 in controlling the robustness of phyllotactic pattern formation.

We initially identified DNE1 in UPF1 IPs, raising the possibility that this protein could be the plant equivalent of the PIN domain endoribonuclease SMG6 in animal (Chicois et al., 2018). We found that, UPF1 co-purifies with DNE1 (44 spectra; Supplementary Table S7) and is also detected with DCP2 (8 spectra; Supplementary Table S6). This observation indicates first that UPF1 is coupled with the decapping complex in plants, as described in mammals (Lejeune et al., 2003) and that it is in addition coupled to a protein complex containing DCP1 and the endonuclease DNE1. A recent analysis of RNA fragments stabilized in *xrn4* reports the accumulation of NMD targets intermediates produced by endonucleolytic cleavage in plants (Nagarajan et al., 2019). Our results put DNE1 in the position of being the favored candidate to exert such an activity.

An interesting parallel can be made between our findings and one of the results of the recent systematic analysis by *in vivo* proximity-dependent biotinylation (BioID) of 119 human proteins associated to mRNA biology (Youn et al., 2018). This analysis identified correlated patterns between endogenous preys and enabled the definition of 144 core components of SGs and P-bodies. Among the P-body components, a correlation cluster appeared around decapping factors (cluster2). Interestingly, this cluster contained the closest homolog of DNE1 in human, the endonuclease MARF1. In both cases, MARF1 and DNE1 represent NYN domain proteins very tightly associated with proteins of the decapping complex. An in-depth molecular phylogenic study across phyla is necessary to reveal whether both genes have a common ancestry, originate from an early horizontal gene transfer or are the results of convergent evolution between evolutionary distant phyla.

Our study reveals a synergistic phyllotactic phenotype observed in *dne1 dcp2*, which can lead to two alternative models for DNE1 action. Either DNE1 acts together with the decapping complex, in a specific complex containing DNE1-DCP1-DCP2 working in parallel of the canonical DCP1-DCP2 decapping complex, or DNE1 exists in an alternative DNE1-DCP1 complex sharing targets with the decapping complex. The first possibility is reminiscent of the mode of action of its human homologue MARF1, proposed to recruit decapping factors to stimulate the degradation of its target (Nishimura et al., 2018). We do not have experimental data supporting the existence of a DCP1-DCP2-DNE1 complex. Our study of proteins associated with DCP1, DCP2 and DNE1 is more in favor of the second hypothesis and suggests the existence in plants, of a protein complex based on DCP1-DNE1 in parallel of the canonical decapping complex based on DCP1-DCP2. Comparing DCP1, DCP2 and DNE1 protein copies per cell (pcc) as found in a recent global proteomic analysis (Bassal et al., 2020) reveal that DCP1 is the more abundant protein (234128 pcc) followed by DCP2 (67345 pcc) and finally DNE1 (791 pcc). This data supports the existence of a main complex containing DCP2 and of an accessory complex containing DNE1. The importance of this accessory complex is revealed by its compensatory effect upon the partial loss of DCP2 function in the weak *dcp2*^*its1*^ mutant, demonstrating its importance for phyllotactic pattern formation. A functional link between DNE1 and DCP2 actions is further supported by our observation that the stable overexpression of DNE1 leads to transcriptomic changes related to the one observed in decapping mutants and to an increased accumulation of targets of the decapping complex (Fig. 7). One way among many to construe this observation is that DNE1 overexpression could compromise the formation of the decapping complex by competing for a common partner, like DCP1. However, a detailed study of the interaction domains between DNE1 and DCP1 will be required in order to test this hypothesis experimentally.

Interestingly, the phyllotactic defects observed in *dne1 dcp2*^*its*^ are also detected in *xrn4* mutant, impaired in 5’-3’ RNA degradation (Fig. 8). This observation supports a possible genetic redundancy between DCP2 and DNE1 in mRNA degradation. This genetic redundancy also provides a plausible explanation to the absence of deregulated genes in *dne1* single mutant, DNE1 as part of an accessory complex, becoming limiting only in case of DCP2 deficiency. Phyllotaxis is controlled by an intricate hormonal balance involving both auxin and cytokinin (Reinhardt et al., 2003; Besnard et al., 2014). Further studies will be required to decipher the RNA substrates targeted by DNE1, DCP2 and XRN4 explaining this observation. Identifying DNE1 targets may also benefit from a recent study indicating that the OST-HTH domains of both DNE1 and its homologue MARF1 have affinity for RNA G-quadruplex (G4), a very specific RNA tertiary structure (Ding et al., 2020). These data suggest that both DNE1 and MARF1 can target G4 containing RNAs and open very interesting perspectives for the study of the targets and biological functions of these NYN domain endoribonucleases.

## MATERIAL AND METHODS

### Plant material

Plant lines used: *dne1-1* (Salk_132521); *vcs-6* (SAIL_831_D08); pDCP1::YFP-DCP1 in *dcp1-3* (SAIL_377_B10); p35S::GFP-DCP2 in *tdt-1* (a *dcp2* mutant); pPABP2::tRFP-PABP2 in WT Col, p35S::GFP-DNE1 in *dne1-1*, p35S::GFP-DNE1-D153N in *dne1-1;* p35S::RFP-DNE1 in *dne1-1; dne1-2* and *dne1-3* were created by the CRISPR*/Cas9* system using the sgRNA target sequence: TCTTCAGGACGTACATCGCT inserted in pKIR1.1 to target DNE1 NYN domain (Tsutsui and Higashiyama, 2017). For IP experiments plants are grown on soil in long-day light conditions (16/8) until flowering and unopened flower buds are collected. For RNAseq experiments surface sterilized seeds (70% ethanol, 0.1% Triton-X 100, washed 5 min with 100% ethanol) are sown on solid MS medium (MS0255 Duchefa®, 1% agar, 1% sucrose, pH 5.7). After 48h of stratification at 4 °C seedlings are grown in long-day light conditions at 21 °C for 14 days.

### Plasmids

Sequence from genomic DNA has been cloned in pB7WGF2, pH7WGR2 gateway destination vectors harboring GFP, RFP sequences respectively to produce N-ter tagged versions of DNE1 both wild-type and catalytic mutant D153N. DNE1 and DCP1 sequences amplified from cDNA were cloned into pGBT9 and pGADT7 to perform Y2H.

### Immunopurifications

Details about samples and replicates of co-immunopurification (IP) experiments are provided in Supplementary Table S1. For IPs 0.3 g of flower buds were ground in 1.5 ml of ice-cold lysis buffer (50 mM Tris-HCl pH 8, 50 mM NaCl, 1 % Triton X-100, supplemented with Roche cOmplete, EDTA-free Protease Inhibitor Cocktail). After cell debris removal by centrifugation (twice 10 min at 16,000 g, 4°C), supernatants were incubated for 30 min with 50 μl of magnetic microbeads coupled to anti-GFP antibodies (Miltenyi). Beads were loaded on magnetized μMACS separation columns equilibrated with lysis buffer and washed four times with 200 μl of washing buffer (20 mM Tris-HCl pH 7.5, 0.1 % Triton X-100). Samples were eluted in 100 μl of pre-warmed elution buffer (50 mM Tris-HCl pH 6.8, 50 mM DTT, 1 % SDS, 1 mM EDTA, 0.005 % bromophenol blue, 10 % glycerol). Negative control IPs were performed under the exact same conditions with WT plants. For IPs crosslink, 0.3 g of flower buds were ground during 10 min in 2.25 ml of ice-cold lysis buffer supplemented with 0.375 % formaldehyde (Thermo Fisher Scientific). The crosslinking reaction was quenched by adding glycine at a final concentration of 200 mM for 5 min. After cell debris removal by centrifugation (twice 15 min at 10,000 g, 4°C), supernatants were incubated for 45 min with 50 μl of magnetic microbeads coupled to anti-GFP antibodies (Miltenyi). Beads magnetic capture and washing steps were done according to the manufacturer’s instructions, except that washes were performed with 50 mM Tris-HCl pH 7.5, 50 mM NaCl, 0.1% Triton X-100, supplemented with Roche cOmplete, EDTA-free Protease Inhibitor Cocktail. Samples were eluted as previously described. Negative control IPs were performed with beads coupled to anti-GFP antibodies in WT plants.

### Mass spectrometry analysis and data processing

Eluted proteins were digested with sequencing-grade trypsin (Promega) and analyzed by nano LCMS/MS on a QExactive + mass spectrometer coupled to an EASY-nanoLC-1000 (Thermo Fisher Scientific). IP data were searched against the TAIR 10 database with a decoy strategy. Peptides were identified with Mascot algorithm (version 2.5, Matrix Science) and data were imported into Proline 1.4 software. IP data were searched against the TAIR 10 database with a decoy strategy. Peptides were identified with Mascot algorithm (version 2.5, Matrix Science) and data were imported into Proline 1.4 software (Bouyssié et al., 2020). Proteins were validated on Mascot pretty rank equal to 1, Mascot score above 25, and 1% FDR on both peptide spectrum matches (PSM score) and protein sets (Protein Set score). The total number of MS/MS fragmentation spectra was used to quantify each protein from at least three independent IPs. Volcano plots display the adjusted p-values and fold changes in Y and X-axis, respectively, and show the enrichment of proteins co-purified with tagged proteins as compared to control IPs. The statistical analysis based on spectral counts was performed using a homemade R package (https://github.com/hzuber67/IPinquiry4) that calculates fold change and p-values using the quasi-likelihood negative binomial generalized log-linear model implemented in the edgeR package. The size factor used to scale samples were calculated according to the DESeq2 normalization method (i.e., median of ratios method). P-value were adjusted using Benjamini Hochberg method from stats R package.

### Phylogeny and structural modlisation

Phylogenetic trees were done with the http://www.phylogeny.fr web tool. Multiple alignments have been made using MUSCLE (v3.8.31). Positions with gaps have been eliminated. The phylogenetic tree has been built using the maximum likelihood method implemented in PhyML (v3.1/3.0 aLRT, approximate likelihood-ratio test). Bootstrap values were calculated on 100 iterations. Graphic representation of the tree was done with TreeDyn (v198.3). The structures of AtDNE1 NYN and OST-HTH domains were modelized using Phyre2: Protein Homology/analogYRecognition Engine V 2.0 (Kelley et al., 2015). The alignment of the structural models produced with NYN HsMARF (PDB ID: 6fdl) and OST-HTH MmMARF1 (PDB ID: 5yad) were done using PyMOL software.

### Agroinfiltration

Competent agrobacteria (GV3101 pMP90) were transformed with vectors containing fluorescent fusion proteins DNE1 WT, mutant D153N, GFP reporter gene or the silencing suppressor P14. *N. benthamiana* leaves were co-infiltrated with bacterial suspension containing DNE1 WT, or D153N, reporter GFP and P14 in a 1:1:1 ratio (Garcia et al., 2014). Infiltrated tissues were collected 3 days after infiltration.

### Subcellular localization analysis

Seven day-old epidermal roots cells of stable *A. thaliana* lines expressing the desired constructs were imaged using a LSM780 confocal microscope (Zeiss) with a 40X objective. Co-localization analysis was performed with ImageJ as follows: foci in images were determined with a user-supervised local maxima detection method (script available on demand). Local intensities in channels visualizing GFP or RFP fusion proteins were measured for every detected focus and the reported values were then charted in a (IGFP versus IRFP) scatter plot for further qualitative assessment of fluorescent spot content correlation.

### Libraries preparation and RNA seq analysis

Transcriptomic analysis was performed on biological triplicates of 14 day-old seedlings of the *dne1-1* mutant, two independent DNE1 OE lines analyzed together (P35S-GFP::DNE1), an DNE1_D153N OE line (35S:GFP-DNE1_D153N). Purified total RNAs were quantified by Qubit (Invitrogen) fluorimeter, RNA’s quality was tested using Bioanalyzer 2100 (Agilent) system. Two micrograms of RNAs were used for libraries preparation with the mRNA-Seq Library Prep Kit V2 for Illumina Platforms (Lexogen) using manufacturer’s instructions. Libraries were sequenced by single read (50 cycles) at GenomEast sequencing platform (IGBMC, Strasbourg) with an Illumina HiSeq 4000. Fastq files were generated using RTA (v2.7.3) and bcl2fatq (v2.17.1.14). Sequences were mapped on *A. thaliana* genome (TAIR10) with Hisat2 (v2.1.0) using default parameters except for intron lengths that have been reduced at 2000 bp. After alignment, the number of reads mapped to each gene were counted with HTseqcount (v0.10.0) using the annotation document Araport11_GFF3_genes_transposons.201606.gff. Statistical analysis was performed on R (v3.6) with DEseq2 package (v1.24.0) and its implemented negative binomial distribution. Adjusted p-values were calculated using the Benjamini and Hochberg method.

### Statistical analysis

Statistical analyses were performed using R 3.6.1, Rstudio 1.2 and the following R packages: DEseq2 1.24.0, stats 3.6.1, multcompView 0.1-8, Limma 3.40.6. For all analyses, a p-value of 0.05 was defined as the threshold of significance. For each figure, the exact value of n and the test used for the statistical analysis are indicated in the figure or in the corresponding legend. Fold change and p-values in Fig. 1b, 1c, 2b and 3 were computed using the quasi-likelihood negative binomial generalized log-linear model implemented in the edgeR package. Correlation coefficients and their associated p-value shown in Fig. 4 were calculated using Pearson correlation method. Statistical significance shown in Fig. 7E and 7G were obtained using Pairwise Wilcoxon Rank Sum Tests with data considered as paired and unpaired, respectively. In Fig. 7G, the proportion of uncapped RNA was retrieved from Anderson et al. 2018 (GSE108852) for all transcripts detected in both this dataset and our RNA experiment. The proportion of uncapped RNA were measured in Col0 and corresponds to the ratio of the RPM from GMUCT for a given transcript (2 biological replicates) normalized by the RPM for that same transcript (4 biological replicates). Boxplot analysis were then performed to compare the proportion of uncapped RNA in all transcripts, non-regulated transcripts, and in transcript either up or down-regulated in D153N transgenic lines and *vcs* mutant. The identification of statistically significant overlap between gene-expression signatures in Venn diagrams shown in Fig. 7 are provided in TableS9 and were computed using the hypergeometric distribution implemented in the stats R package (v3.6.1). Statistical significance shown in Supplementary Fig S5 were obtained using the Limma moderated F-test. All p-value were adjusted using the Benjamini Hochberg method.

### Quantitative real-time reverse transcriptase PCR (qRT-PCR)

Three to six independent biological replicates of 14 days old seedlings were analyzed (see Supplementary Fig S5). RNA was extracted using TRI Reagent® (Sigma), genomic DNA was removed using the DNase RQ1 (PROMEGA), cDNA was synthesized using a mix of random hexamers and oligo d(T) by Superscript IV (Invitrogen). Technical triplicates qRT-PCR were performed with SYBR-green I Master Mix (Roche) using Light Cycler 480 (Roche), following the manufacturer’s instructions. mRNA abundance was compared to two reference genes *ACT2* and *TIP41* (primer used in Table S12). The ΔΔCt method was used to calculate relative RNA abundance.

### Phyllotactic pattern measurements

Phyllotactic pattern was assessed on 35 successive siliques per plant on the main stem, starting from the first silique, on fully grown stems. Divergence angles were measured using a dedicated device as previously described (Peaucelle et al 2007). Angles were measured between the insertion points of two successive floral pedicels, independently of the outgrowth direction of the pedicel. Phyllotaxy orientation for each individual, was set to the direction giving the smallest average divergence angle. Measurements were performed on between three and four biological replicates, analysis of individual replicates are shown in Supplementary Fig S6.

### Accession numbers

Sequence data from this article can be found in the GenBank data libraries under accession numbers At1g15560 (DNE1); At1g08370 (DCP1); At5g13570 (DCP2); At3g13300 (VCS); AT3G13290 (VCR); At2g45810 (RH6); At4g00660 (RH8); At3g61240 (RH12); AT5G47010 (UPF1) At3g58570 (RH52); At4g34110 (PABP2); Niben101Scf05368g03015 (NbRBC); Niben101Scf00508g00007 (NbPLAST); Niben101Scf11490g00008 (NbTHIAM).

### Large datasets

RNAseq datasets generated during this study have been deposited in NCBI’s Gene Expression Omnibus and are accessible through GEO Series accession number GSE155806.

Mass spectrometry proteomics raw data have been deposited to the ProteomeXchange Consortium via the PRIDE partner repository with the dataset identifier n° PXD020780, reviewer access: /Za2LUVLe.

Data shown in Supplementary Figure S1 were extracted from the pastDB project at http://pastdb.crg.eu/; (Martín et al., 2021).

## SUPPLEMENTAL DATA

The following supplemental materials are available.

**Supplementary Figure S1**. Graphical representation showing normalized RNA-seq data for the expression of *DNE1, DCP1* and *DCP2* in *Arabidopsis* at 90 different developmental stages.

**Supplementary Figure S2**. Accumulation levels of RFP-DNE1 and RFP-D153N in the transient expression assay.

**Supplementary Figure S3**. Characterization of GFP-DNE1 transgenic lines.

**Supplementary Figure S4**. Characterization of GFP-D153N transgenic lines.

**Supplementary Figure S5**. Q-PCR validations of transcriptomic data on selected deregulated genes.

**Supplementary Figure S6**. Density plots showing the quantification of divergent angles for each biological replicate separately.

**Supplementary Table S1**. Details of biological material used for IPs.

**Supplementary Table S2**. List of proteins significantly enriched in YFP-DCP1 IPs.

**Supplementary Table S3**. List of proteins significantly enriched in GFP-DCP2 IPs.

**Supplementary Table S4**. List of proteins significantly enriched in GFP-DNE1 IPs.

**Supplementary Table S5**. List of proteins significantly enriched in crosslinked YFP-DCP1 IPs.

**Supplementary Table S6**. List of proteins significantly enriched in crosslinked GFP-DCP2 IPs.

**Supplementary Table S7**. List of proteins significantly enriched in crosslinked GFP-DNE1 IPs.

**Supplementary Table S8**. Amino acid identity and similarity of DNE1 protein domains (NYN, OST1, OST2) from representative streptophytes species.

**Supplementary Table S9**. Comparison between lists of differentially expressed genes using hypergeometric test.

**Supplementary Table S10**. Statistical analysis of the Q-PCR validations by moderated t-test.

**Supplementary Table S11**. Aligned sequences from *dne1-2* and *dne1-3* CRISPR-Cas9 mutants.

**Supplementary Table S12**. Analysis of the distribution of divergent angles in different biological replicates using the Kolmogorov-Smirnov test.

**Supplementary Table S13**. DNA oligonucleotides used in this study.

## ACKNOWLEDGEMENTS

The authors thank C. Bousquet-Antonelli and J.M. Deragon for pDCP1::YFP-DCP1, pPABP2::tRFP-PABP2 constructs and lines. This research was funded by the Centre National de la Recherche Scientifique (CNRS) and performed in the frame of the Interdisciplinary Thematic Institute IMCBio, as part of the ITI 2021-2028 program of the University of Strasbourg, CNRS and Inserm, was supported by IdEx Unistra (ANR-10-IDEX-0002), by SFRI-STRAT’US project (ANR 20-SFRI-0012), and EUR IMCBio (IMCBio ANR-17-EURE-0023) under the framework of the French Investments for the Future Program » as well as from the previous LabEx NetRNA (ANR-10-LABX-0036). The authors also acknowledge the funding of a QExactive Plus mass spectrometer (ThermoFisher) by an IdEx grant from the University of Strasbourg. The funders had no role in study design, data collection and analysis, decision to publish, or preparation of the manuscript.

## FOOTNOTES

This research was funded by an attractivity grant from the NetRNA LabEx, ANR-10-LABX-0036_NETRNA (DG).

Da.G. conceived the original research plans, supervised the work, analyzed the data and wrote the manuscript with major contributions of Do.G. and M.S.; M.S. performed and analyzed the transcriptomic and mass spectrometry experiments with help from P.H., J.C. L.K and HZ; C.C. performed preliminary experiments and colocalization studies; H.Z. developed scripts using R and data analysis; A.G. performed the phylogenetic analysis and 3D modeling of DNE1; A.P performed northern blot analyses; T.C. performed phenotypic analysis; E.U. performed western blot analyses; J.M. assisted with microscopy and built the system to measure phyllotaxis. Da.G. agrees to serve as the author responsible for contact and ensures communication.

